# Distinct postsynaptic morphogenetic strategies across *Drosophila* embryonic muscles during neuromuscular junction formation

**DOI:** 10.64898/2026.03.20.713228

**Authors:** Melissa Ana Inal, Daichi Kamiyama

**Affiliations:** Department of Cellular Biology, University of Georgia, Athens, GA, USA; Neuroscience Program, University of Georgia, Athens, GA, USA

**Keywords:** Synaptic partner selection, Neuromuscular junction, *Drosophila*, Myopodia, Muscle-specific GAL4 drivers, Embryonic development, Cell–cell recognition

## Abstract

Precise synaptic connectivity emerges through coordinated interactions between neurons and their target cells during development. At the Drosophila embryonic neuromuscular junction (NMJ), postsynaptic muscle fibers actively participate in this process by extending dynamic, actin-rich protrusions termed myopodia that interact with approaching motor growth cones.

Previous work focusing on muscle 12 (M12) revealed that myopodia cluster at nascent neuron–muscle contact sites, suggesting that specialized postsynaptic architectures may facilitate synaptic partner selection. However, whether similar morphogenetic strategies operate across the diverse set of embryonic muscles has remained unclear. Here, we establish a genetic imaging toolkit that enables minimally invasive visualization of defined muscle subsets throughout the embryo. Using muscle-specific and stochastic GAL4 drivers to label muscle membranes *in vivo*, we systematically compare myopodial organization across multiple muscle fibers, including M12, M14, M6, and M7. We find that postsynaptic morphology varies substantially between muscles. M12 displays robust myopodial clustering associated with a prominent sheet-like membrane structure, which we term the muscle lamella, whereas M6 and M14 frequently form myopodial clusters but do not evidently exhibit this structure. In contrast, M7 shows markedly reduced clustering frequency and smaller clusters. These observations reveal previously unrecognized heterogeneity in postsynaptic organization among neighboring muscles during early neuromuscular development. Together, our findings demonstrate that myopodial clustering represents a broadly deployed but differentially organized strategy by which muscles engage motor axons during synaptic partner selection. The imaging toolkit established here provides a foundation for systematic analysis of neuron–muscle interactions across the embryonic musculature and reveals that distinct muscles employ diverse morphogenetic strategies during NMJ assembly.

## INTRODUCTION

Model organisms have been indispensable for uncovering the mechanisms that govern synaptic partner selection. The fruit fly *Drosophila melanogaster* has emerged as a particularly powerful system owing to its sophisticated genetic toolkit, compact and tractable nervous system, and rapid developmental timeline. Because cell–cell interactions unfold with exceptional spatial and temporal precision, live imaging is critical for visualizing the dynamic steps of circuit formation. Recent advances in deep-tissue imaging have enabled direct observation of olfactory receptor neuron axons and projection neuron dendrites as they target specific glomeruli, revealing highly dynamic axonal and dendritic behaviors that sculpt glomerular development [1, 2].

The *Drosophila* neuromuscular junction (NMJ) provides a well-established model for investigating synaptic partner selection [3, 4]. Its defined anatomy, dynamic cellular interactions, and the superficial positioning of muscles in transparent embryos make it ideally suited for high-resolution live imaging [5, 6]. Within each abdominal hemisegment, 33 motor neurons innervate a stereotyped set of 30 muscle fibers, generating a reproducible connectivity map (**Fig. 1**) that remains largely conserved through larval stages [7, 8]. This remarkable precision has made the embryonic NMJ an invaluable platform for dissecting the cellular events that govern synaptic specificity.

**Fig. 1.**
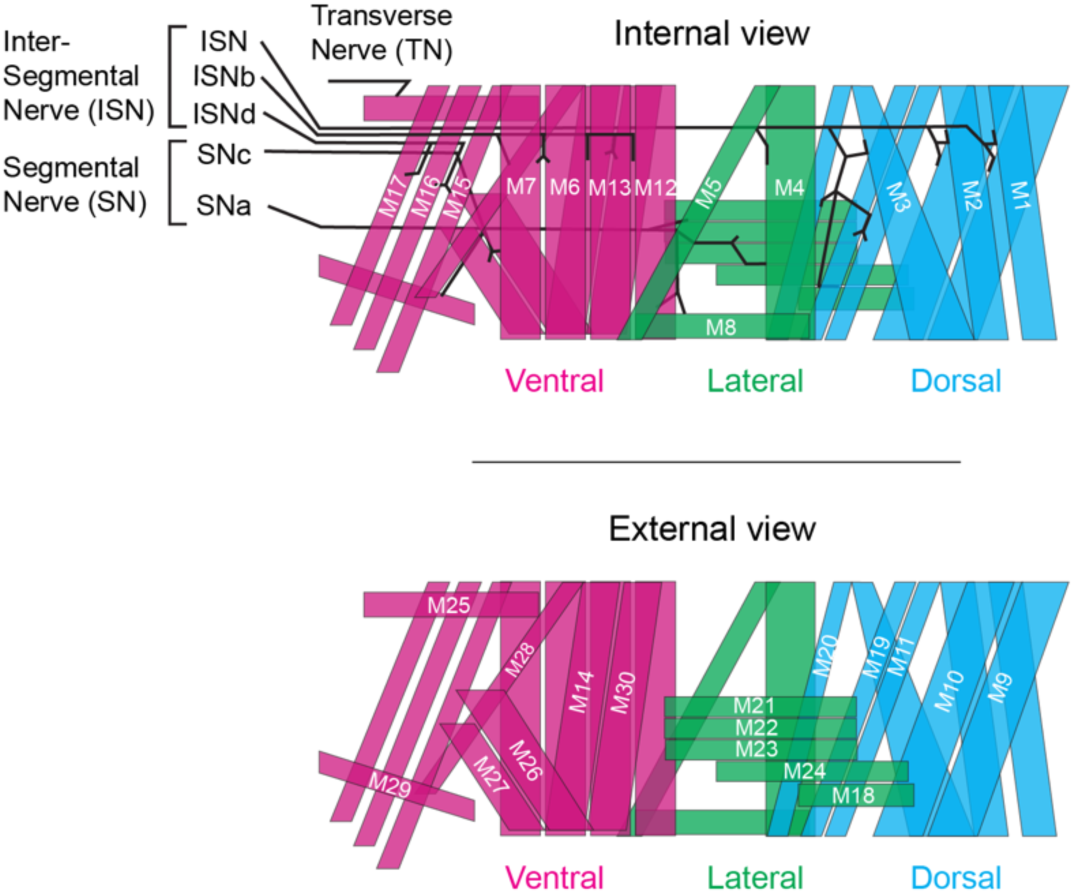
Internal and external schematic views of embryonic body wall musculature. Internal and external schematic views of musculature. **Top:** Internal view of musculature with the innervation pattern of each nerve. Nerve tracks are labeled on the left side of the schematic. Only the internal muscles are labeled. **Bottom:** External view of the musculature where external muscles are labeled separately from the internal muscles. Musculature is colored based on the defined groupings in the main text: ventral in magenta, lateral in green, and dorsal in cyan.

As motor axons exit the ventral nerve cord (VNC), they extend into well-defined territories and survey their environment to identify appropriate muscle targets [9-14]. Early models proposed that growth cone filopodia were the primary structures responsible for sampling muscle-derived cues. This view shifted with the discovery of dynamic filopodia on postsynaptic muscle cells—termed myopodia by Ritzenthaler et al.—first identified at the embryonic NMJ [15] and later observed in embryonic mouse muscle and in cultured rodent and Xenopus muscle cells [16-19]. Myopodia are highly motile protrusions capable of extending, retracting, and directly engaging with axonal filopodia, positioning them as active contributors to the partner-matching process.

Much of our current understanding of myopodial dynamics derives from studies of muscle 12 (M12) and its corresponding motor neurons [15, 20, 21]. This system benefits from a single-muscle-specific driver (*tey*-*GAL4*) [15, 22] that, together with membrane-targeted GFP, enables selective labeling of the M12 membrane and high-resolution visualization of postsynaptic behavior during partner selection. Myopodia form clusters that contact approaching motor neuron growth cones [15, 20]. Electron microscopy has further revealed that axonal filopodia and myopodia interdigitate at nascent synaptic sites, providing ultrastructural evidence that myopodia serve as a postsynaptic counterpart to axonal exploration [15, 23].

Genetic evidence reinforces this model. Kohsaka et al. demonstrated that the leucine-rich repeat transmembrane protein Capricious (Caps), expressed as a GFP fusion in M12 using *tey*-*GAL4*, localizes to the tips of myopodia where initial contacts with axonal filopodia occur [20]. In *caps* mutants—or in double mutants lacking both Caps and its paralog Tartan—the number of initial contacts between M12 and its presynaptic growth cones is markedly reduced.

Moreover, the number of nascent synaptic sites on M12 is diminished at later stages. These findings indicate that myopodia rely on tip-localized molecular cues to detect and engage axonal filopodia, contributing directly to synaptic partner matching.

Whether the structural transitions and molecular strategies identified in M12 extend broadly across the embryonic musculature remains unresolved. Previous efforts to visualize additional muscle fibers, such as M6, using live fillet-dissected embryos and lipophilic dyes (DiI) detected myopodial contacts in approximately 46% of hemisegments at 13 hr AEL [24]. This lower detection rate—compared with the near-complete visualization achieved for M12 using *tey-GAL4*—may reflect technical limitations, including phototoxicity during DiI-based live imaging that can induce rapid filopodial retraction. Alternatively, intrinsic biological differences among muscle fibers may influence myopodial stability, abundance, or morphology, and certain muscles may employ distinct strategies for synaptic partner selection. Therefore, systematic and minimally invasive imaging approaches are required to determine whether the mechanisms defined in M12 represent a generalizable feature of neuromuscular development.

To address these challenges, we developed a genetic toolkit by screening GAL4 drivers expressed in defined subsets of muscle fibers. This collection includes seven GAL4 lines that stably label distinct muscle groups and three additional drivers that enable stochastic single-cell labeling across the musculature. Together, these lines support expression of transgenes—including membrane-targeted fluorescent proteins—for imaging live or fixed muscle tissues with high spatial resolution. Using this approach, we confirmed that M12 muscles form clustered myopodia at high frequency (∼90% of hemisegments) and identified a sheet-like membrane structure at the base of these protrusions, which we term the muscle lamella. Importantly, we found that M6—previously examined using more invasive approaches—and M14, which had not been characterized at this resolution, also generate myopodial clusters at presumptive synaptic sites with comparable frequency. In contrast, M7 forms myopodial clusters less frequently and with smaller cluster size. Notably, the muscle lamella was observed only in M12, whereas the other muscles examined did not display this structure. These findings reveal variability in how individual muscle fibers deploy myopodial remodeling during early neuromuscular development.

Together, this toolkit enables systematic analysis of postsynaptic contributions to synaptic partner selection across the embryonic musculature and reveals that myopodial clustering is not uniformly deployed across muscles. Instead, distinct muscle fibers appear to utilize different morphological strategies during neuromuscular partner matching.

## RESULTS

### Anatomical Organization of Embryonic Muscles and Innervation

We first outline the anatomical framework of the embryonic neuromuscular system. We follow the standard nomenclature widely used in the field, in which muscles are identified by their anatomical position and designated by number [3, 7, 8, 14, 25]. This organizational context provides a foundation for interpreting the expression patterns described in later sections.

Embryonic muscles receive innervation from motor neurons projecting through two primary nerve branches: the segmental nerve (SN) and the intersegmental nerve (ISN) (**Fig. 1**). A notable exception is the transverse nerve (TN), which courses along the segmental boundary to innervate M25. Subdivisions of the SN and ISN further refine neuromuscular connectivity, segregating inputs to ventral, lateral, and dorsal muscle groups.

The ventral muscle group is innervated by SNc (segmental nerve c), ISNd (intersegmental nerve d), and ISNb (intersegmental nerve b). This group includes the ventral longitudinal muscles (M12, M13, M6, and M7); the ventral oblique muscles (M14, M30, M28, M15, M16, and M17); and the ventral acute muscles (M26, M27, and M29) (**Fig. 1**; magenta).

The lateral muscle group is innervated primarily by SNa (segmental nerve branch a), which targets the lateral transverse muscles (M21, M22, M23, and M24); the lateral oblique muscle (M5); and the segmental border muscle (M8). This group also includes the dorsal transverse muscle (M18) and the lateral longitudinal muscle (M4), which, although positioned laterally, receive innervation from the ISN (**Fig. 1**; green).

The dorsal muscle group is innervated predominantly by the ISN and comprises the dorsal acute muscles (M1, M2, and M3) and the dorsal oblique muscles (M9, M10, M11, M19, and M20) (**Fig. 1**; cyan).

Together, these categories highlight the strong correspondence between muscle position and neuronal input, an important consideration for evaluating the specificity of GAL4 drivers in our screening strategy.

### Screening Strategy to Identify GAL4 Lines Specific to Subsets or Single Muscles

To date, only a small number of GAL4 drivers—such as *tey-GAL4*—have been reported to selectively label discrete subsets of embryonic muscle fibers. To expand the toolkit of muscle-selective drivers, we screened an extensive collection of enhancer-driven GAL4 lines generated at the Janelia Research Campus. Although these lines were originally developed to assay genomic enhancer activity across mostly neuronal cell types [26], their expression patterns in embryonic body wall musculature remain incompletely characterized. Using the associated imaging datasets, we established a systematic and unbiased screening pipeline.

As summarized in **Fig. S1**, we began by reviewing images of stage 16 (13–16 hr AEL) embryos with annotated body wall musculature across 2,480 GAL4 lines available through the FlyLight collection at the Janelia Research Campus. Lines exhibiting restricted expression in small subsets of muscles were prioritized, whereas those showing broad epithelial or ubiquitous expression were excluded, yielding a refined pool of 85 candidates. These candidates were subsequently crossed to a membrane-targeted GFP reporter (*UAS-gapGFP*) and screened in dissected embryos imaged at 10× magnification using a confocal microscope.

This secondary screen identified 10 promising lines, which we grouped into two categories based on expression consistency. The first category consisted of lines that reproducibly labeled stable, anatomically coherent clusters of muscles across multiple segments and embryos. In contrast, several lines displayed variable expression patterns, labeling different subsets of muscles either between embryos or between adjacent segments within the same embryo. Such variability likely reflects incomplete or fragmented enhancer regions [27]. While consistently expressed lines are particularly valuable for studying muscle groups as coordinated units, variably expressed drivers offer unique advantages for achieving single-muscle resolution. Accordingly, these variable-expression lines formed the second category.

### Identification and Characterization of Seven GAL4 Drivers with Stable Expression in Small Subsets of Embryonic Muscles

We identified seven GAL4 driver lines that reproducibly label discrete subsets of embryonic body wall muscles (**Fig. 2**). Across abdominal segments A2–A6, their expression patterns showed no detectable differences along the anteroposterior axis. Within individual segments, signal intensity varied slightly among labeled muscle fibers (**Fig. S2**), but overall labeling patterns remained highly consistent throughout the 11–16 hr AEL developmental window.

**Fig. 2.**
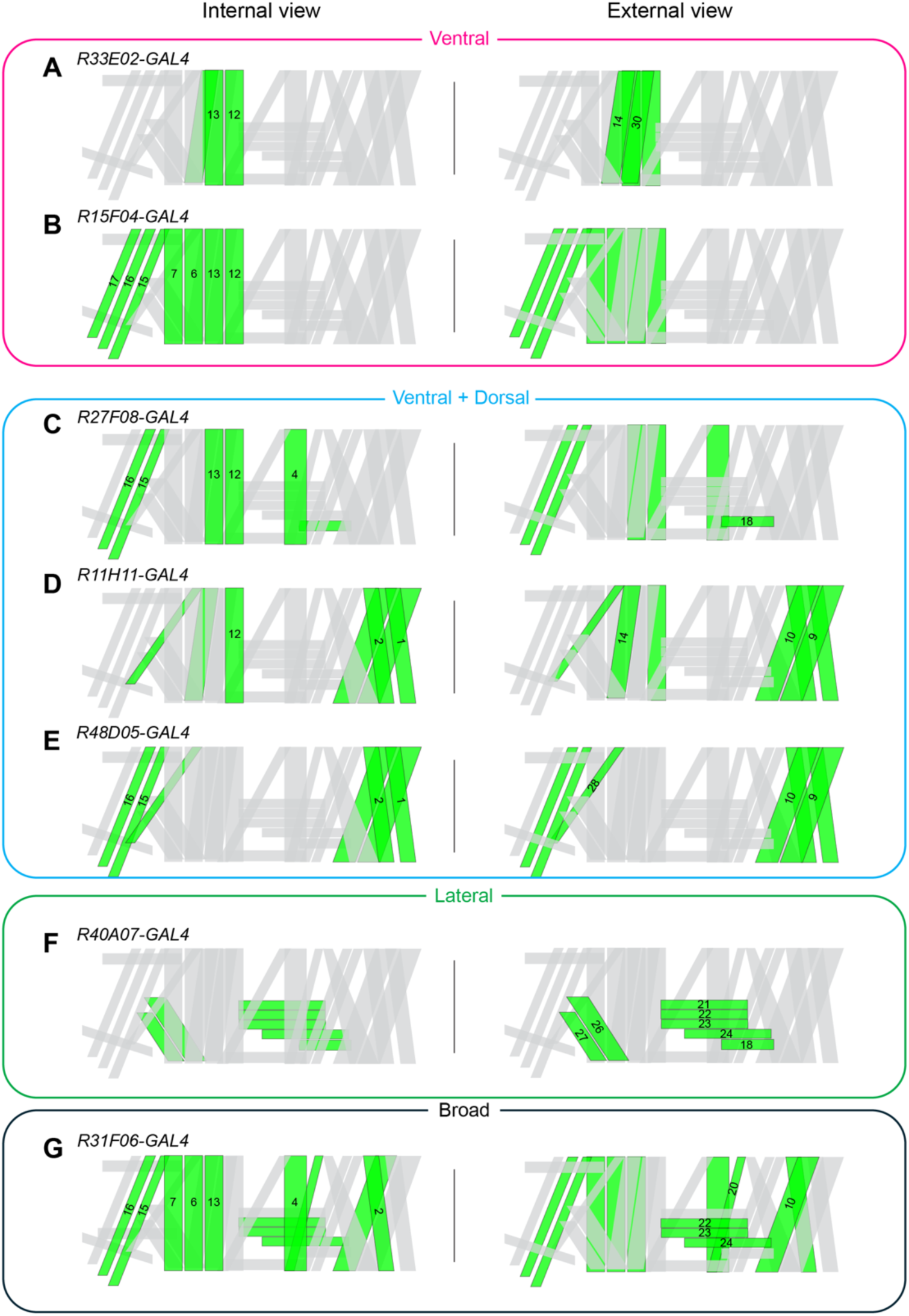
Muscle-specific stable GAL4 drivers. GAL4 drivers with distinct spatial expression patterns in embryonic muscles are shown. (**A**, **B**) Ventral muscle activity, (**C**–**E**) combined ventral and dorsal muscle activity, (**F**) lateral muscle activity, and (**G**) broad muscle activity across multiple groups. Raw data are shown in Fig. S2.

Based on their spatial expression profiles, these drivers were grouped into four categories: ventral (three lines), lateral (one line), ventral/dorsal (two lines), and broad (one line labeling muscles across multiple positional domains).

The ventral category includes two drivers with strong, selective expression in ventral musculature. *R33E02-GAL4* primarily labels M14 and M30 and also marks ventral longitudinal muscles M12 and M13 (**Fig. 2A**). *R15F04-GAL4* provides the broadest ventral coverage, labeling M12, M13, M6, and M7, as well as M15, M16, and M17 (**Fig. 2B**). Together, these lines serve as precise tools for examining ISNb-innervated ventral muscles.

Two additional drivers show expression spanning both ventral and dorsal domains. *R27F08-GAL4* labels M15, M16, M12, and M13, together with dorsal muscles M4 and M18 (**Fig. 2C**). R11H11 and R48D05 share similar dorsal expression patterns—M1, M2, M9, and M10—but differ in their ventral profiles. *R11H11-GAL4* displays broader ventral activity, labeling M15, M16, M17, M12, and M6 (**Fig. 2D**). In contrast, *R48D05-GAL4* is more selective, labeling M28, M15, M16, and M17 (**Fig. 2E**). These drivers provide complementary tools for comparative analyses of coordinated ventral–dorsal subsets and support studies of innervation by ISN, ISNb, and ISNd.

The lateral category is represented by *R40A07-GAL4*, which predominantly labels external musculature. This driver marks the lateral transverse muscles M21, M22, M23, and M24, the dorsal transverse muscle M18, and the ventral external muscles M26 and M27 (**Fig. 2F**). *R40A07-GAL4* provides a valuable tool for investigating SNa innervation of transverse muscles and SNc innervation of ventral external muscles.

Finally, *R31F06-GAL4* displays selective yet widespread expression across ventral, lateral, and dorsal domains while maintaining near single-muscle resolution. In the ventral region, it labels M14, M30, M15, M16, M17, and M12. In the lateral region, M22, M23, M24, and M4 are marked. In the dorsal region, M1, M2, M9, and M10 are labeled (**Fig. 2G**). This versatile driver supports integrated analyses across anatomical regions and facilitates cross-domain comparisons.

Together, these seven GAL4 lines provide robust and reproducible labeling of distinct muscle subsets, offering reliable tools for dissecting the organization and developmental patterning of the embryonic body wall musculature.

### GAL4 Expression in Random Subsets of Muscles for Single-Muscle Studies

Our second set of GAL4 drivers was selected for their variable yet informative expression patterns across segments within individual embryos. This variability produces low-frequency but highly valuable labeling events, enabling single-muscle resolution across a broad range of muscle types. We focused on three lines—*R9H06-GAL4*, *R23B04-GAL4*, and *R28E05-GAL4*—each exhibiting medium to high overall coverage while maintaining sufficient stochasticity to target individual muscles.

*R9H06-GAL4* displays relatively unbiased and frequent stochastic activity, making it one of the most effective lines for single-muscle analyses. Variability commonly manifests as segment-specific differences within the same embryo (**Fig. 3A**). In our assessment of 36 hemisegments from five embryos, this line labeled 29 of the 30 muscles, with M29 being the only muscle not observed. Some muscles, however, were labeled at comparatively low frequencies: in the ventral region, M27 and M25 appeared in fewer than 10% of observations; in the lateral region, M21 and M24 were rarely labeled; and in the dorsal region, M20 showed the lowest frequency (**Fig. 3B**).

**Fig. 3.**
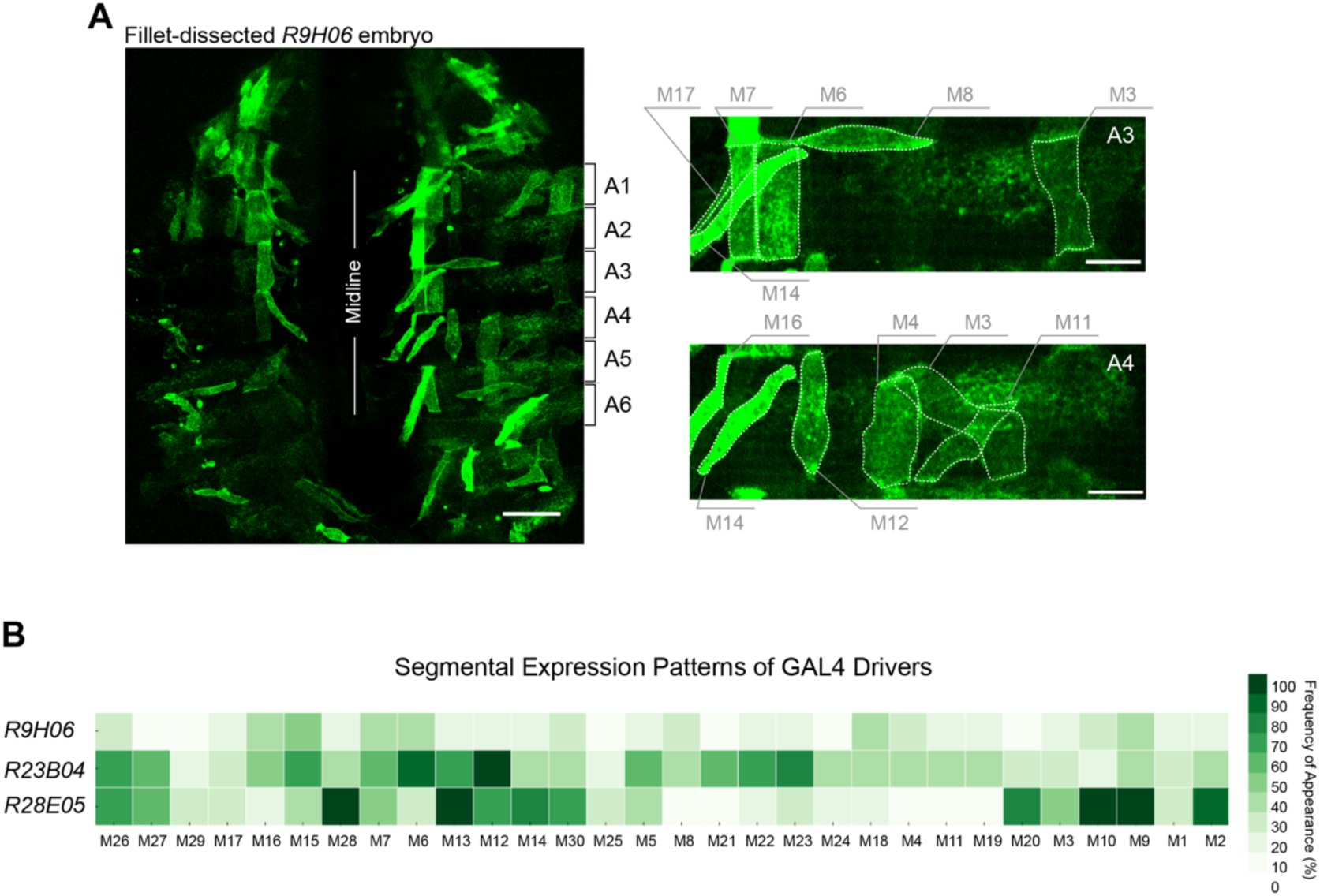
GAL4 drivers with stochastic muscle activity. **(A)** *R9H06-GAL4* crossed to *UAS-CD4::tdGFP* was filleted as an embryo at 13 hr AEL and immuno-stained with anti-GFP. Two panels are magnified right abdominal hemisegments, A3 and A4, with muscle fibers outlined from the complete embryo demonstrating the variation of expression among segments. Some fluorescence signal comes from the epithelial cells. Scale bars: 50 µm (left); 20 µm (right). (**B**) Expression frequency of each muscle between A1-A6 segments for the GAL4 drivers (*R9H06*, *R23B04*, and *R28E05*) that have stochastic single cell activity. The layout is organized from ventral to distal domains of body wall; each cell has a muscle ID indicated below the heatmap. *n* = 36, 30, and 30 hemisegments (5 embryos each), respectively.

*R23B04-GAL4* labeled some muscles at single-muscle resolution, including M26, M27, M5, M8, M21, M22, M23, and M24 (**Fig. 3B** and **S3A**). This driver was particularly valuable for targeting M29, which was not labeled by *R9H06-GAL4*. Additional ventral longitudinal muscles were also labeled: M7, M13, and M12 appeared stochastically, whereas M6 was consistently labeled across segments. Notably, M15, M16, and M17 near the VNC exhibited combinatorial and segment-specific expression patterns.

*R28E05-GAL4* labeled several muscles that were underrepresented in the other stochastic lines, showing low-frequency expression in M13, M14, and M20 (**Fig. 3B** and **S3B**). This line displayed broad ventral and dorsal coverage but lacked expression in the lateral transverse muscles. Its complementary expression pattern relative to the other stochastic drivers enables near-comprehensive coverage of the muscle field when used in combination.

Together, these three drivers provided at least 20% labeling frequency for every muscle, including lower-frequency muscles such as M25, M29, and M17. Most muscles exhibited labeling frequencies of 30% or higher. Although only *R9H06-GAL4* was used for the experiments described in the following section, the choice of driver ultimately depends on the experimental context and the desired muscle coverage for reproducible single-muscle resolution.

### Myopodial Clustering During Early Innervation in M12 and M14

Myopodia are fine, actin-rich protrusions that extend from muscle fibers during early stages of neuromuscular development [15, 28]. Owing to their slender and highly dynamic structure, live imaging is particularly well suited for accurately capturing their native morphology and behavior *in vivo*. This principle was first demonstrated in 2000 [15], when the Chiba laboratory used membrane-targeted GFP driven by *tey-GAL4* to selectively label the plasma membrane of M12, enabling continuous visualization of myopodial activity throughout development.

Prior to 13 hr AEL, myopodia are broadly distributed across the surface of M12. As development progresses into the 15–16 hr AEL window, these protrusions gradually consolidate into a smaller, well-defined region of the muscle membrane. This phenomenon—known as myopodial clustering—consistently occurs at the site where M12 normally receives innervation from the ISNb motor axon (V and RP5 motor neurons [3, 4]; see also **Fig. 4A**).

**Fig. 4.**
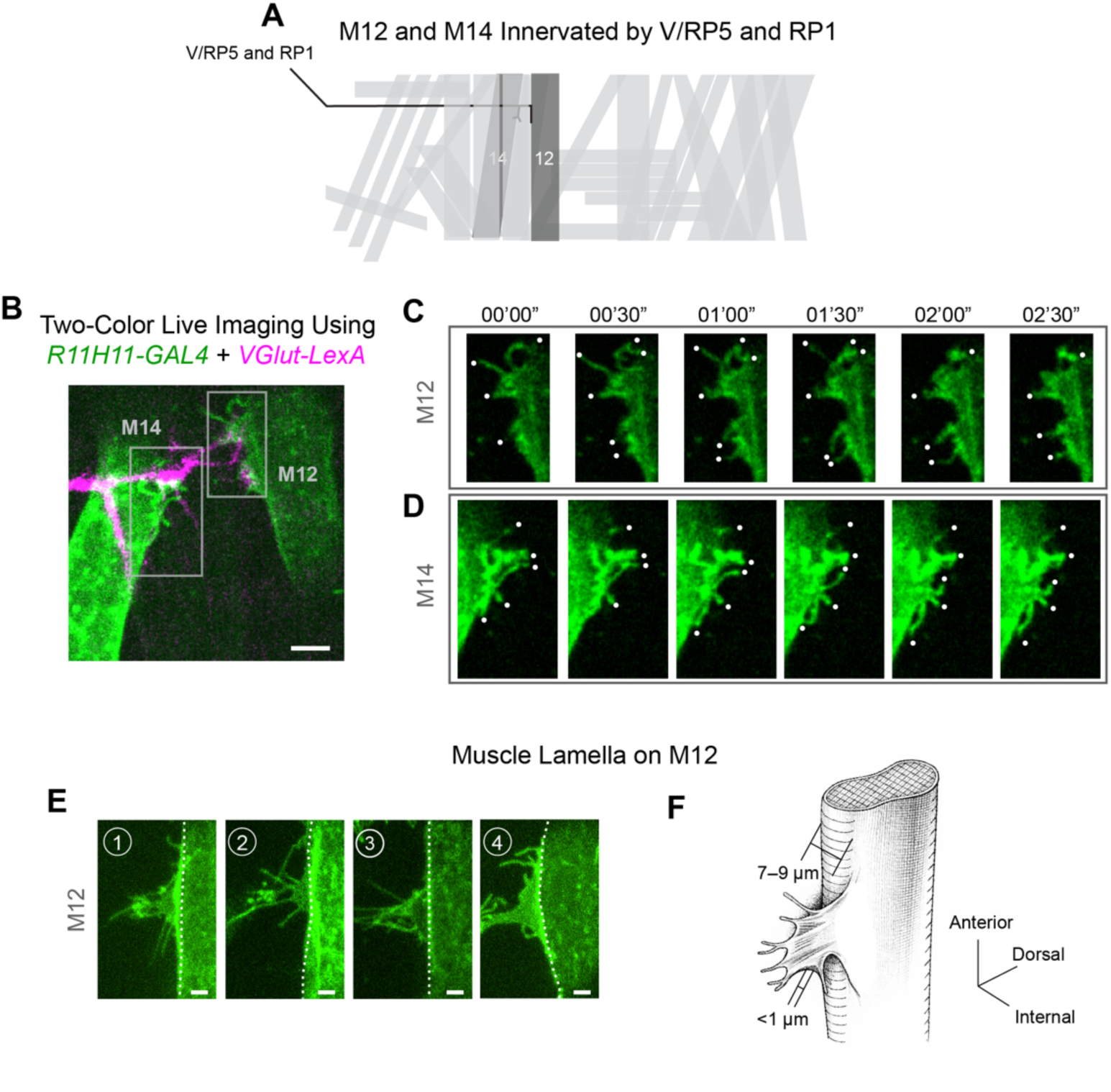
Live visualization of M12 and M14 muscle fibers during early neuron–muscle contact. **(A)** Schematic illustration of M12 and M14 innervated by the V/RP5 and RP1 motor neurons, respectively [3, 4]. (**B**) Neurons and muscles were visualized in dissected embryos using *VGlut-LexA* driving *LexAop2-CD4::tdTomato* and *R11H11-GAL4* driving *UAS-CD4::tdGFP*, respectively. Neuron–muscle contact sites on M12 and M14 were imaged at 15.5 hr AEL. The ventral nerve cord (VNC) is positioned on the left, and motor nerves extend from left to right toward their target muscles. The gray boxes indicate axon branch contact points with M12 and M14, respectively. Scale bar: 5 µm. (**C**–**D**) Live imaging of M12 and M14 at the neuron–muscle contact sites indicated by the upper and lower gray boxes in panel B. Still frames are taken from Movies 1 and 2, corresponding to panels C and D. (**E**) Four examples of M12 from independent embryos showing the sheet-like membrane protrusion (muscle lamella). White dotted lines indicate the boundary between the main M12 muscle fiber edge and the sheet-like membrane extension. Scale bars: 2 µm. (**F**) Schematic illustration of the muscle lamella based on our observations.

To demonstrate comparable live, single-muscle imaging using our toolkit, we used *R11H11*-*GAL4*, which provides reliable single-cell resolution in M12. For two-color membrane visualization, this driver was paired with an orthogonal LexA/LexAop system, using a motor neuron–specific *VGlut* gene-trap LexA line [29]. Embryos expressed *UAS*-*CD4::tdGFP* in muscle and *LexAop2*-*CD4::tdTomato* in motor neurons, enabling simultaneous visualization of both membranes (**Fig. 4B**).

We focused on the developmental stage when M12 forms stable contacts with the ISNb branch (15.5 hr AEL). At this stage, M12 engages ISNb axons at the presumptive synaptic site (**Fig. 4C**). Consistent with earlier descriptions, dynamic filopodia remain clearly visible, forming a localized cluster (**Fig. 4C** and **Movie 1**). During live imaging, we also observed robust filopodial activity in the neighboring muscle fiber M14, which is GAL4-positive in *R11H11*-*GAL4*. Filopodia on M14 were concentrated in the anterodorsal region of the muscle (**Fig. 4D** and **Movie 2**). Across both muscle fibers, filopodial clustering occurred in more than 90% of hemisegments (90.5% in M12, *n* = 85; 90.2% in M14, *n* = 41). These findings suggest that myopodial remodeling is not restricted to a single muscle but may occur broadly across multiple muscle fibers during early innervation.

Inspection of time-lapse recordings also revealed a sheet-like membrane structure on M12, which we refer to as the muscle lamella (see also Discussion). The morphology of the muscle lamella varies among M12 fibers (**Fig. 4E**). When projected into two dimensions, the area of the lamellar region is typically 14.4 ± 1.5 µm^2^ (n = 6 lamellae). This structure is extremely thin (typically <1 µm in thickness) compared with the overall muscle fiber diameter (approximately 7–9 µm) (**Fig. 4F**). Notably, the lamella is positioned along the internal face of the muscle fiber adjacent to the developing contact site.

We define a lamella as a continuous sheet-like membrane extension larger than 5 µm^2^. This threshold was chosen because smaller protrusions are difficult to distinguish confidently from densely packed filopodia near the muscle edge at the spatial resolution of our fluorescence imaging. Using this criterion, we did not detect lamellae in M14 at this developmental stage (*n* = 41; data not shown). Although smaller sheet-like membrane expansions were occasionally observed in M14, they remained below this size threshold, suggesting that while similar protrusive remodeling may occur in both muscles, the extent of membrane spreading differs substantially between M12 and M14.

### Myopodial Clustering on M7 and M6

In addition to M12, earlier work by Ritzenthaler et al. reported myopodial clustering on M6; however, the frequency of this event was below 50% in the segments examined [24]. At that time, no GAL4 driver was available for selective labeling of M6, and clustering was assessed using DiI-based membrane labeling in live, fillet-dissected embryos at 13 hr AEL. Although dye labeling provides strong membrane contrast, lipophilic dyes can induce substantial phototoxicity, making this approach poorly suited for extended live imaging [30]. Consequently, the original method may have reduced detection efficiency by promoting rapid retraction or loss of delicate myopodia.

To reassess this question, we performed fillet dissections comparable to those used previously but rapidly fixed the preparations immediately after dissection. Rapid fixation with freshly prepared, EM-grade paraformaldehyde has been recently shown to better preserve fine filopodial structures while minimizing structural collapse [31]. This strategy also enabled subsequent immunofluorescence labeling using anti-HRP (horseradish peroxidase), a pan-neuronal membrane marker that reliably outlines motor axons. In contrast, *VGlut-LexA* labeling used in earlier sections of this study is not sufficiently robust at 13 hr AEL because expression at this stage remains relatively weak. Anti-HRP staining therefore allowed reliable identification of presumptive neuron–muscle contact sites at this developmental time point.

To label M6, we employed the stochastic driver *R9H06-GAL4* in combination with *UAS-CD4::tdGFP*, which marks this muscle at moderate frequency with single-muscle resolution. Using this approach, we quantified myopodial clustering at 13 hr AEL, when the RP3 motor neuron innervates the M7/6 cleft (**Fig. 5A**). Specifically, we asked whether M6 forms myopodial clusters near this presumptive contact zone. In embryos in which M6 was labeled, we observed clustering of M6 filopodia, with frequencies exceeding 88.4% (*n* = 43; see also **Fig. 5B**), substantially higher than previously reported frequency of 46% [24]. For consistency, we used the same definition of myopodial clustering as in the previous study (more than three myopodia at the presumptive contact zone; see Methods for details). These results demonstrate that M6 robustly forms myopodial clusters at sites of early motor contact.

**Fig. 5.**
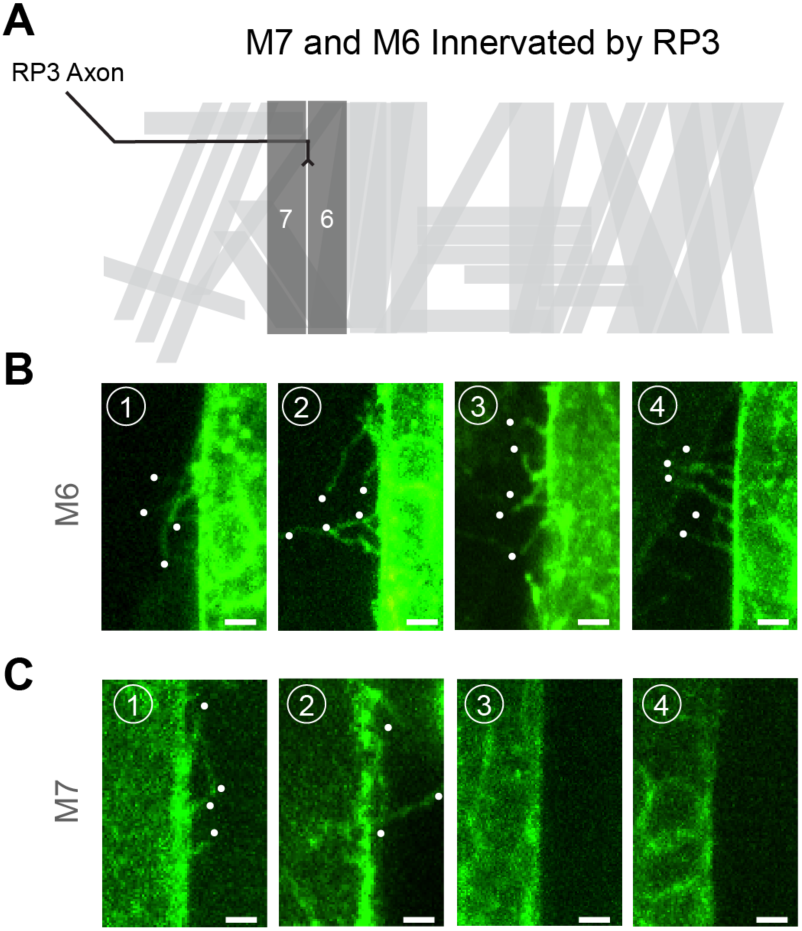
Visualization of M7 and M6 muscle fibers and myopodial clustering. **(A)** Schematic illustration of M7 and M6 innervated by the RP3 motor neuron. (**B**–**C**) Myopodia were visualized using the stochastic driver *R9H06-GAL4* in combination with *UAS-CD4::tdGFP*. Embryos were immunostained with anti-GFP and anti-HRP antibodies. Myopodial clusters are shown at presumptive neuron–muscle contact zones, identified by HRP staining of motor axons contacting the muscle fibers (images not shown). Representative images of M6 and M7 muscle fibers at 13 hr AEL are shown from four independently selected contact sites in panels (**B**) and (**C**), respectively. Scale bars: 2 µm.

Given the close anatomical relationship between M7 and M6, we next quantified clustering on M7 near the RP3 contact site (**Fig. 5A**). Initial analysis revealed a lower clustering frequency than that observed for M6 (40.4%, *n* = 47; Fisher’s exact test, *p* = 3 × 10^-6^; see also **Fig. 5C**). In addition, the number of filopodia on M7 appeared to be reduced by approximately 50% compared with those observed on M6 within a myopodial cluster. However, M7 labeling was less bright (**Fig. 5B–C**), potentially limiting detection sensitivity and reducing the clarity of individual filopodia. To address this concern independently of genetic labeling, we performed DiD labeling (a spectrally distinct lipophilic dye related to DiI) after fixation to eliminate phototoxicity-associated artifacts while enhancing membrane contrast. This approach allowed clear visualization of individual myopodia (**Fig. S4**). Importantly, clustering remained absent in a substantial fraction of hemisegments, with frequencies comparable to those obtained using the genetically encoded approach (44%, *n* = 25; Fisher’s exact test, *p* = 0.81). These findings indicate that the reduced penetrance reflects intrinsic biological variability rather than a technical limitation.

Together, these results establish myopodial clustering as a reproducible and biologically regulated feature of early neuromuscular development. Clusters consistently emerged at presumptive contact sites in multiple muscle fibers, supporting a conserved role for myopodial remodeling in postsynaptic partner selection. In contrast to M12, the other muscles examined did not display a detectable muscle lamella at the corresponding developmental stages (M6 and M7 at 13 hr AEL; M12 and M14 at 15.5 hr AEL). We also observed substantially lower clustering frequency and smaller clusters in M7, suggesting the presence of muscle-specific developmental programs that differentially regulate or modulate clustering during synaptic partner matching.

## DISCUSSION

Our analysis of stable and stochastic GAL4 drivers expands the experimental toolkit for visualizing NMJ development at single-muscle-fiber resolution. The resulting expression patterns indicate that myopodial clustering—previously emphasized in M12—is more broadly distributed across the embryonic musculature than recognized previously. Multiple fibers, including M6, M7, and M14, exhibit focal zones of dense myopodia that frequently coincide with motor axon approach or early neuron–muscle interactions. Notably, clustering is not uniform across fibers, underscoring the importance of examining multiple muscles to capture the full spectrum of developmental strategies.

### Muscle-Specific Regulation of Myopodial Clustering and Synaptic Partner Matching

A central question is whether most embryonic muscle fibers intrinsically possess the capacity to form myopodial clusters or whether this ability is restricted to subsets defined by molecular identity, spatial position, or developmental timing. In the muscles examined here, all fibers exhibited myopodial clustering; however, clear differences were observed in cluster morphology, density, and frequency across muscles. Our analyses to date have focused primarily on ISNb motor axons and several of their partner muscle fibers, including M6, M7, M14, and M12.

The situation in M7 is particularly informative. M7 and M6 are unique in that a single motor neuron, RP3, innervates both adjacent fibers and later forms synapses within their shared cleft. Despite this common presynaptic partner and close anatomical proximity, M6 displayed highly penetrant clustering, whereas M7 showed reduced and more variable clustering. How this discrepancy influences synaptic partner matching and subsequent synaptogenesis remains unresolved. Notably, a preliminary SEM analysis by Yoshihara et al. [32] reported that at the varicosity-forming stage (approximately 19 hr AEL), both large and small varicosities are present in the region between the two fibers, with a greater number associated with M6 than M7. These observations raise the possibility that early differences in myopodial clustering may correlate with later asymmetries in synaptic maturation, although this relationship will require further experimental validation.

A similar anatomical arrangement exists between M14 and M30, which are both innervated by RP1 motor neurons [3, 4]. While M14 exhibits prominent and readily detectable filopodial clustering (**Fig. 4**), it remains unclear whether M30 engages the clustering program to a comparable extent or instead exhibits reduced clustering, similar to the relationship between M7 and M6. Systematic analysis of this muscle pair will provide an additional test of muscle-specific regulation of early postsynaptic remodeling.

Together, these comparisons raise broader questions about how motor neurons select synaptic partners when multiple potential targets lie in close proximity. In several cases, adjacent muscles share a synaptic cleft, requiring a single motor axon to determine whether to stabilize contacts with one or both muscles. For example, RP3 ultimately forms synapses with both M6 and M7 within their shared cleft, and RP1 similarly innervates both M14 and M30. In contrast, V/RP5 motor neurons form synapses with M12 but not with the neighboring M13 despite their close spatial proximity. One possibility is that differences in myopodial clustering across muscles provide an early morphological indicator of how muscles participate in synaptic partner selection. Variations in protrusive activity or clustering frequency could influence whether a motor growth cone stabilizes contacts with one muscle, both muscles, or neither within a shared synaptic environment.

### Temporal Control of Myopodial Cluster Formation

Defining when myopodial clusters first emerge is essential for understanding their functional role. Previous studies guided our initial imaging windows—approximately 13 hr AEL for M7 and M6 and 15.5 hr for M12 [15, 28]—but these time points likely represent snapshots within a broader and potentially fiber-specific temporal program.

The mechanisms underlying cluster formation remain incompletely defined. Two general models may explain the relationship between cluster formation and axon arrival. In one model, clusters arise before axonal contact and function as prepatterned landmarks that guide growth cone approach. In an alternative model, clusters assemble or stabilize only after neuron–muscle interactions begin. Previous observations in M12 strongly support the latter model: in *prospero* mutants, in which motor axon outgrowth is halted, myopodial clusters fail to form [15]. In addition, studies in rodent culture systems demonstrate that neuron-derived agrin can induce myopodia-like protrusions [19]. Similarly, whole-mount analysis of the embryonic mouse triangularis sterni muscle has shown that neurotransmitter signaling (ACh) can destabilize excess myopodial clusters [16]. Together, these studies suggest that neuronal signals directly trigger and/or stabilize muscle protrusive activity through local diffusible or contact-mediated mechanisms.

Current evidence therefore supports a model in which cluster formation occurs within a developmental window aligned with motor axon entry into the body wall and the initiation of target sampling. Axon arrival and synaptic maturation proceed along a ventral-to-dorsal gradient [32], raising the possibility that ventral fibers initiate clustering earlier, whereas dorsal fibers do so later. Defining the temporal relationship between axon arrival and cluster emergence across diverse muscle types will be essential for understanding how spatial and temporal cues are integrated to establish precise neuromuscular connectivity.

### A Muscle Lamella Forms at the Developing M12 Neuromuscular Junction

We observed that M12 muscle fibers form a distinct sheet-like membrane structure (**Fig. 4E–F**). The original study describing myopodia briefly mentioned a similar feature [15], referring to it as a “lamellipodium” or “lamellipodia-like structure.” Canonical lamellipodia are broad membrane protrusions supported by a branched actin network and are typically found at the leading edge of migrating cells and axonal growth cones, where they exhibit rapid and highly dynamic movements [33-35]. In contrast, the sheet-like structure observed on M12 displays markedly different properties. It forms a thin membrane sheet but extends slowly along the internal surface of the muscle fiber toward the ventral side over a period of approximately 30 minutes (M.A.I. and D.K., unpublished observations). The cytoskeletal organization underlying this structure remains unknown, and its dynamics differ substantially from those of classical lamellipodia. Based on its morphology, position, and distinctive behavior, we therefore redefine this structure as the muscle lamella, a thin membrane sheet that emerges at the neuron–muscle contact interface during early neuromuscular development.

Notably, we observed this structure only in M12 under the conditions examined here. M12 is unique among the muscles analyzed in that it receives input from the neuromodulatory V motor neuron, which forms peptidergic type III boutons containing dense-core vesicles loaded with insulin-like peptides [8, 36]. One possibility is that development of this specialized synapse, such as the type III bouton, requires the presence of the muscle lamella. Alternatively, similar sheet-like structures may occur more broadly at other neuromuscular junctions but only transiently or during narrowly defined developmental windows that were not captured in our current imaging paradigm.

### Molecular Mechanisms of Muscle–Axon Recognition

The molecular cues that mediate early muscle–axon recognition remain a major open question. Capricious (Caps), an immunoglobulin superfamily protein implicated in M12 targeting, remains a compelling candidate [20, 37]. However, its restricted expression—limited to muscles M1, M2, M9, M10, M12, M15, M16, and M17—indicates that additional factors must contribute to the widespread clustering observed across the musculature. A broader set of cell-surface proteins (CSPs) has been identified as potential recognition molecules [37-45]. Whether these proteins localize specifically to myopodial tips, as observed when Caps is overexpressed using *tey-GAL4*, and whether such localization is required for stabilizing early contacts remain important unresolved questions.

A commonly accepted concept is that recognition mechanisms are combinatorial, relying on coordinated use of multiple CSPs rather than a single molecule per muscle [25, 46-48]. Individual fibers may depend on distinct repertoires of adhesion molecules or integrate partially redundant pathways to achieve robust partner matching. Advances in single-cell transcriptomics now enable gene expression profiling at the level of individual embryonic muscle fibers [49]. Integrating these datasets with functional assays using our GAL4-based toolkit will be essential for identifying candidate match-making molecules and elucidating how they collectively shape early neuron–muscle communication.

### Summary and Future Directions

Taken together, these findings support a model in which myopodial clustering represents a widespread, conserved, and actively regulated strategy for establishing precise neuromuscular connectivity. Defining the spatial patterns, temporal dynamics, and molecular cues that govern cluster formation will be essential for developing a unified framework for synaptic partner selection. In addition, we define the sheet-like membrane structure from which clusters emerge as the muscle lamella. The physiological role of this structure remains unclear and will require further investigation.

Future work integrating advanced live imaging, membrane labeling, and endogenous protein tagging [6, 50-53] will further clarify how muscles and motor neurons engage in coordinated communication during circuit assembly. Importantly, the GAL4 drivers characterized here are active early and persist throughout embryogenesis, providing a platform not only for studying synaptic partner matching but also for examining earlier processes such as myoblast fusion and later stages of synapse establishment and maintenance. This expanded toolkit therefore offers broad opportunities to dissect multiple phases of neuromuscular development within a unified and experimentally tractable framework.

## MATERIALS AND METHODS

### Drosophila lines

All GAL4 lines and LexA lines identified in this study are listed in Table S1. Reporter lines were: *5x UAS-GAP::GFP* (gifted by Carlos Lois) was used for screening GAL4 activity, *UAS-CD4::tdGFP* (RRID: BDSC_35836) was used for higher magnification characterization of the stochastic GAL4 drivers and live imaging experiments. *LexAop2-CD4::tdTomato* (RRID: BDSC_77178) was used for imaging *VGlut-LexA*. Flies were reared at 25°C, and staging have been determined at this temperature based on hours after egg laying and morphological criteria described by Campos-Ortega and Hartenstein in 1997.

### Dissection and immunohistochemistry

Embryonic dissections and immunostaining were performed as previously described. For initial screening of GAL4 drivers, fillet-dissected stage 16 (13 hr AEL) embryos were fixed in 4% paraformaldehyde (Electron Microscopy Sciences) for 5 minutes at room temperature and counterstained with anti-HRP conjugated to Cy3 (1:100; Jackson ImmunoResearch, RRID: AB_2338959) to visualize motor axons. For characterization of stochastic GAL4 drivers crossed to a GFP reporter line, additional immunostaining was performed. Following fixation, samples were washed in PBS containing 0.01% Triton X-100 (PBST) and blocked for 1 hour at room temperature in PBST supplemented with 0.06% BSA. Primary antibodies were diluted in blocking solution, and embryos were incubated overnight at 4°C with monoclonal rabbit anti-GFP (1:200; Thermo Fisher Scientific, RRID: AB_2536526). The following day, samples were incubated for 2 hours at room temperature with secondary antibodies, including donkey anti-rabbit Alexa Fluor 488 (1:200; Thermo Fisher Scientific, RRID: AB_2535792) and anti-HRP Cy3 (1:200). After immunostaining, samples were washed thoroughly and mounted in PBS for imaging.

### Lipophilic dye labeling

DiD (Thermo Fisher Scientifc, Cat# D307) was dissolved in ethanol and vegetable oil to prepare a 5 mg/mL working solution. Dissected embryos were fixed and stained with anti-HRP antibody as described above. The dye was applied directly to the surface of muscle M7 and incubated for 10 minutes. Excess dye was removed by gentle aspiration using a mouth pipette. Samples were then washed and mounted in PBS for imaging. Note that physical access of the injection needle restricts this labeling approach to certain muscle fibers, including M7.

### Confocal microscopy

Confocal microscopy images of fixed, filleted embryos were captured using an inverted fluorescence microscope (Ti-E, Nikon) equipped with 10x 0.30 NA air, 40x 0.80 NA water immersion, or 100x 1.45 NA oil immersion objectives (Nikon). For live imaging, time-lapse movies were acquired using a 40x 1.25 NA silicone immersion objective (Nikon). The microscope was coupled to a Dragonfly spinning disk confocal system (CR-DFLY-501, Andor). Excitation was provided by three solid-state lasers (488 nm, 40 mW; 561 nm, 50 mW; and 642 nm, 110 mW), routed through a multimode fiber and homogenized by the Andor Borealis illumination unit. The imaging path included a Dragonfly laser dichroic (405/488/561/640) and three bandpass emission filters (525/50 nm, 600/50 nm, and 725/40 nm). Images were recorded using an electron-multiplying charge-coupled device (EMCCD) camera (iXon, Andor). All imaging parameters, including laser power, exposure time, and z-step intervals, were optimized for each experiment to minimize photobleaching and ensure consistent signal detection.

### Heatmap generation of GAL4 activity

Each muscle’s expression in single segments (A2–A6) across 5 embryos were counted for each driver. The frequency of expression was split into 10 percent bins. The heatmap was generated using a custom Python script.

### Definition and Quantification of Myopodial Clustering

Myopodia were defined as protrusions extending from muscle fibers with a length of ≥1 µm. A myopodial cluster was defined as the presence of three or more myopodia located within ±1 µm of a presynaptic contact site. Presynaptic terminals were identified using anti-HRP staining or VGlut-LexA–labeled motor neurons, and these markers were used to define neuron–muscle contact zones for quantification.

### Image processing

Images taken using 10x objective were sharpened using the built-in deconvolution algorithm by Fusion (Andor, Oxford Instruments). All images were processed in ImageJ (NIH) to improve brightness and contrast. To smooth for the background artifacts in live movies, a Gaussian blur (radius = 0.5) was applied. Images for stochastic GAL4 drivers were taken using 40x objective and stitched together in ImageJ using the Grid/Collection stitching plugin. Figures were structured using Adobe Illustrator (Adobe).

## ACKNOWLEDGEMENTS

We thank members of the Kamiyama laboratory for helpful discussions and technical assistance, and the Bloomington Drosophila Stock Center for fly stocks. This work was supported by the National Institutes of Health (NIH R21 NS128750 to M.A.I. and D.K.)

## AUTHOR CONTRIBUTIONS

D.K. conceived the study. D.K. and M.A.I. designed the experiments. M.A.I. performed the experiments and analyzed the data. D.K. supervised the project. M.A.I. and D.K. wrote the manuscript.

## DATA AVAILABILITY STATEMENT

All data supporting the findings of this study are available within the paper and its supplementary information files.

